# Human TRPV4 engineering yields an ultrasound-sensitive actuator for sonogenetics

**DOI:** 10.1101/2024.10.16.618766

**Authors:** Lu Zhao, Kevin Xu, Irina Talyzina, Jingyi Shi, Shisheng Li, Yaoheng Yang, Shuming Zhang, Jie Zheng, Alexander I. Sobolevsky, Hong Chen, Jianmin Cui

## Abstract

Sonogenetics offers non-invasive and cell-type specific modulation of cells genetically engineered to express ultrasound-sensitive actuators. Finding an ion channel to serve as sonogenetic actuator it critical for advancing this promising technique. Here, we show that ultrasound can activate human TRP channel hTRPV4. By screening different hTRPV4 variants, we identify a mutation F617L that increases mechano-sensitivity of this channel to ultrasound, while reduces its sensitivity to hypo-osmolarity, elevated temperature, and agonist. This altered sensitivity profile correlates with structural differences in hTRPV4-F617L compared to wild-type channels revealed by our cryo-electron microscopy analysis. We also show that hTRPV4-F617L can serve as a sonogenetic actuator for neuromodulation in freely moving mice. Our findings demonstrate the use of structure-guided mutagenesis to engineer ion channels with tailored properties of ideal sonogenetic actuators.

## Main

The ongoing advancement of neuromodulation techniques has been driven by the pursuit of a profound understanding of brain function and the development of effective treatments for neurological disorders ^1–3^. This advancement necessitates the development of neuromodulation techniques that offer non-invasive and spatiotemporal precise control for research and clinical applications ^4–7^. Current non-genetic neuromodulation techniques, such as deep-brain stimulation, activate many brain circuit components within the targeted region with efficacy that can be difficult to predict or control ^8,9^. Genetic approaches, such as optogenetics and chemogenetics, provide cell- type specifically but suffer from several drawbacks ^10,11^. Optogenetics often requires surgical procedures to deliver light stimuli to targets, particularly those deep in the brain, while chemogenetics lacks temporal resolution. Sonogenetics provides a complementary approach of using ultrasound to control the activities of specific neuronal populations genetically modified to express ultrasound-sensitive ion channels ^12,13^. Among all existing physical energy forms, ultrasound is unique in that it can safely penetrate through the scalp and skull and target any location inside the brain with millimeter-scale spatial resolution ^14–16^. Leveraging the unique capability of ultrasound, sonogenetics offers a non-invasive and spatiotemporally precise approach to modulate specific neural populations throughout the entire brain ^13^. This approach holds immense potential for advancing our understanding of neural circuits and developing precise, non-invasive and precise treatments for neurological disorders.

Sonogenetics requires ultrasound-sensitive ion channels as effective actuators ^13,17^. When propagating through the tissue, ultrasound generates mechanical effects, suggesting that mechanosensitive ion channels can be used as sonogenetic actuators. Since 2016, several mechanosensitive ion channels were shown to be activated by ultrasound, including two-pore domain potassium channels (TREK-1, TREK-2 and TRAAK) ^18,19^, MscL ^20–22^, Piezo1 ^23^, MEC-4 ^24^, prestin ^25^, TRPA1 ^26^, TRPC1, TRPP2, and TRPM4 ^27^. Nevertheless, none of these channels have satisfied all criteria of an ideal sonogenetic actuator: 1) highly sensitive to ultrasound stimulation; 2) precise controllability for activation and deactivation by ultrasound; 3) well-defined ion selectivity; 4) optimal size for easy delivery; 5) non-immunogenicity; 6) non-interference with normal neuronal function; and 7) reduced sensitivity to other stimuli. We therefore embarked on the search for an ultrasound-sensitive ion channel that can be used as an actuator for sonogenetic neuromodulation.

### hTRPV4 is sensitive to ultrasound

The transient receptor potential (TRP) channel TRPV4 is involved ion nociception, thermal-, mechano- and osmo-sensation ^28^ and demonstrates responsiveness to mechanical stress ^29–31^ and osmolarity ^32,33^. We tested whether this mechanosensitive ion channel can serve as an effective sonogenetic actuator and found that human TRPV4 (hTRPV4) can be activated by focused ultrasound (FUS) (**Fig. 1**). Currents through wild-type hTRPV4 (hTRPV4-WT) expressed in *Xenopus* oocytes were elicited by FUS using a 1-MHz transducer and recorded by a two-electrode voltage clamp (**Fig. S1**). Both continued and pulsed FUS (C-FUS and P-FUS) enhanced the currents amplitude, up to 15.07 ± 1.21-fold, depending on the acoustic pressure of FUS (**Fig. 1a, b**) as well as the duration of FUS exposure in P-FUS stimulation (**Fig. S2**). These enhanced currents were due to increased activation of hTRPV4-WT, while could be inhibited by a specific antagonist RN-1734 (**Fig. 1c**). We also demonstrated that FUS activated hTRPV4-WT expressed in mammalian HEK293T cells. Since the channel is Ca^2+^-permeable, Ca^2+^ entry through hTRPV4- WT was visualized with a calcium indicator Fluo-4 AM that was preloaded into the cells. The expression of hTRPV4-WT was monitored by fluorescence of mCherry fused to the channel protein (**Fig. 1d**) and confirmed by the detection of Ca^2+^-influx in response to 4αPDD, a synthetic TRPV4 agonist (**Fig. S3**). Compared to the control non-transfected cells, a notable increase in Ca^2+^ influx was detected in hTRPV4-WT-expressing cells in response to P-FUS stimulation (**Fig. 1d, e**). On average, a 20% increase in the maximum fluorescence intensity (△F/F) was observed in response to P-FUS.

**Fig. 1.**
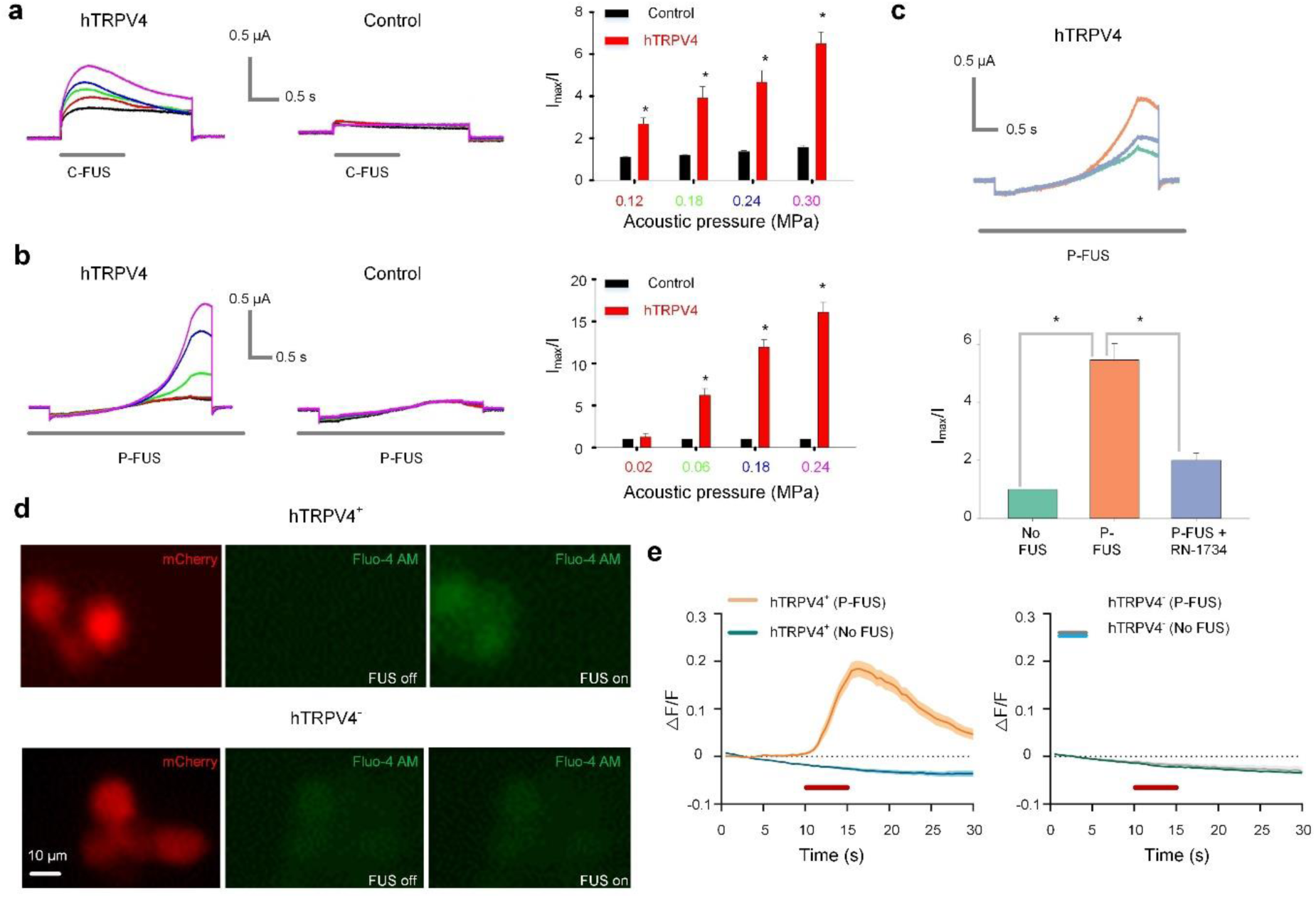
Low-intensity FUS activation of hTRPV4. **a**, C-FUS activation of hTRPV4. Currents of a hTRPV4-expressed oocyte (*left*) and a water-injected control oocyte (*middle*) were elicited by a voltage pulse to +50 mV (**Fig. S2a**). Colors indicate the acoustic pressures of C-FUS as in (*right*), the averaged fold change of current amplitudes elicited by C-FUS at different acoustic pressures. For this and subsequent figures, bars represent the mean ± SEM, *: p < 0.005, n > 3. **b**, P-FUS activation of hTRPV4. Currents were elicited by a -100 mV to +60 mV voltage ramp (**Fig. S2b**). P-FUS had a DC of 20%, PRF of 10 Hz, and various acoustic pressures. **c**, RN-1734 (3 μM) suppression of P-FUS induced hTRPV4 currents. Colors indicate the same in current traces (*upper*) and averaged current amplitudes normalized to the maximum P-FUS induced currents (*lower*). P- FUS: 0.12 MPa, 10% DC, 10 Hz PRF. **d**, Fluorescence images of HEK293T cells transfected with hTRPV4 (plenti-CaMKⅡ-hTRPV4-WT-p2A-mCherry) or control (plenti-CaMKⅡ-p2A-mCherry) plasmids. Red: HEK293T cells expressing mCherry with (*left*, *top*) or without (*right*, *top*) hTRPV4. Green: intracellular Ca^2+^ before and after P-FUS of 1.7-MHz, 0.59 MPa, 40% DC, 10 Hz, 5 s duration. **e**, Normalized fluorescence change (△F/F) in hTRPV4-expressed (*left*) and control (*right*) cells induced by P-FUS or no FUS. Solid lines and shadow indicate the mean ± SEM. Red bars: P-FUS on.

### Mutations alter hTRPV4 sensitivity to ultrasound and other stimuli

The hTRPV4 channel is activated by multiple stimuli, including hypo-osmolarity, elevated temperature, and chemical ligands ^34–37^. Accordingly, the lack of specific sensitivity to ultrasound compromises the use of hTRPV4-WT as a sonogenetic actuator. To overcome this difficulty, we screened nine-point mutations distributed across various structural domains of hTRPV4 (**Fig. S4a**) that were reported to reduce channel sensitivity to elevated temperature, hypo-osmolarity, or agonist 4αPDD ^38,39^. Consistent with the previous reports, our own measurements confirmed that these mutations reduced hTRPV4 sensitivity to one or more of the three stimuli (**Fig. 2a-c, S4d- f**). Interestingly, some of these mutations also altered the channel sensitivity to ultrasound (**Fig. 2d, S4g**), suggesting that this sensitivity is an intrinsic property of hTRPV4 and dependent on the structure of channel protein. Previous studies suggested that hTRPV4 sensitivity to agonist 4αPDD and elevated temperatures may share the same molecular mechanism ^36^. Consistently, we found that the changes in hTRPV4 sensitivity to these stimuli are well correlated (**Fig. 2a, c**). On the other hand, each individual mutation increased or decreased sensitivity of hTRPV4 to ultrasound regardless of changes in sensitivity to other stimuli (**Fig. 2**, **Table S1**). Thus, ultrasound appears to be activating stimulus of hTRPV4 orthogonal to hypo-osmolarity, elevated temperature, and ligands.

**Fig. 2.**
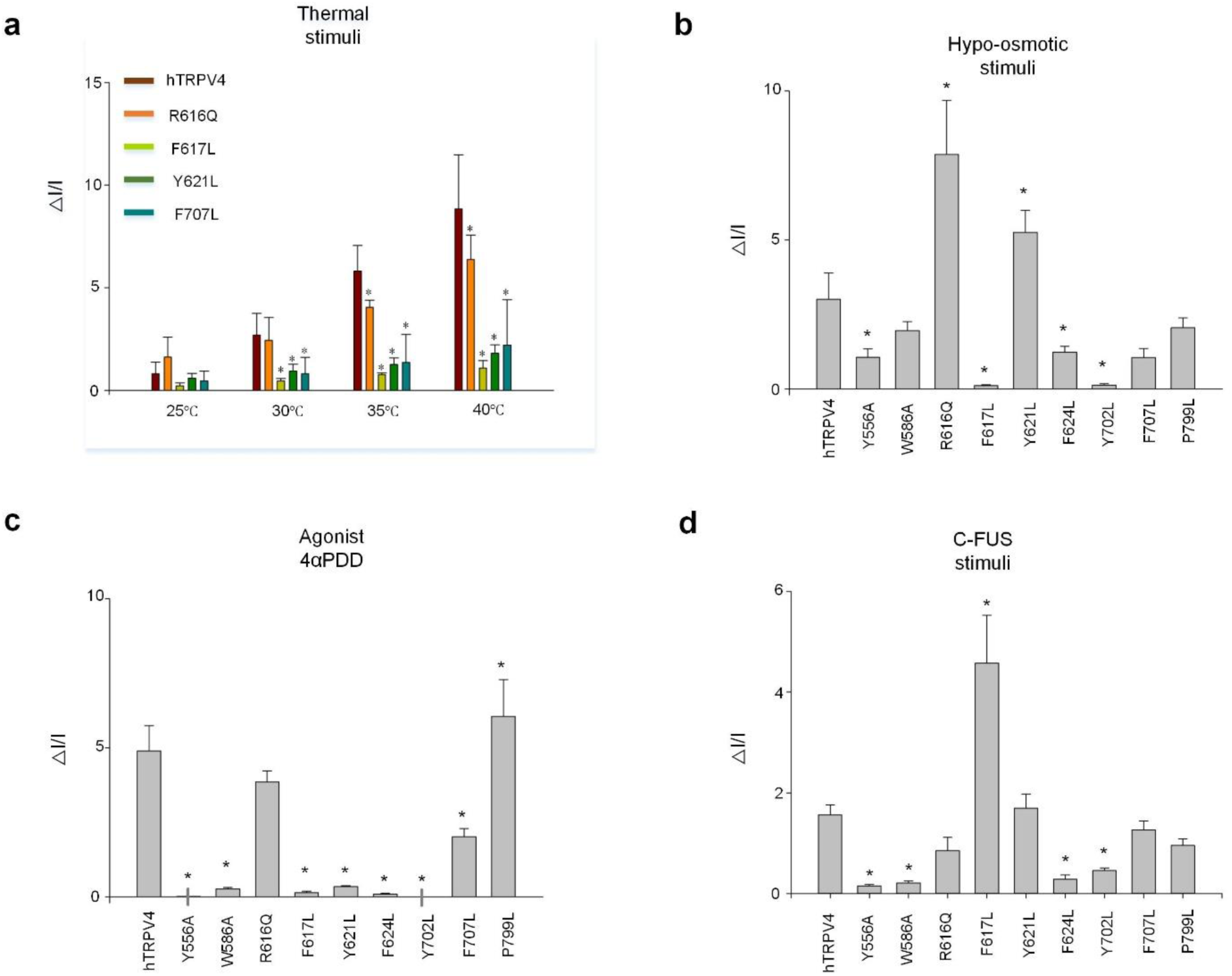
Sensitivity changes of mutant hTRPV4 to ultrasound and other stimuli. **a-d**, Averaged fold change of current amplitudes of the hTRPV4-WT and hTRPV4 mutants activated by temperature (**a**), hypo-osmolarity (**b**), 4αPDD (**c**), and C-FUS (0.18 MPa for 1 s) (**d**). *: mutant results significantly differ from that of WT (p < 0.005, n > 3).

### hTRPV4-F617L is a sonogenetic actuator

One of the characterized mutations, F617L, enhanced sensitivity of hTRPV4 to ultrasound but dramatically reduced sensitivity to elevated temperature, hypo-osmolarity, and agonist 4αPDD (**Fig. 3**). The subsequent experiments demonstrated that F617L enhanced FUS sensitivity of the channel at all acoustic pressures (**Fig. 3a**). On the other hand, at the membrane voltages of -100 mV to +100 mV, which cover the physiological range and beyond, the F617L mutation caused reduction in current elevation in response to hypo-osmolarity (**Fig. 3b**, **S5**) or agonist 4αPDD (**Fig. 3c**). Sensitivity of the channel to elevated temperature was also significantly reduced (**Fig. 3d**). These results show that the mutant hTRPV4-F617L channel is more specifically sensitive to ultrasound stimulation.

**Fig. 3.**
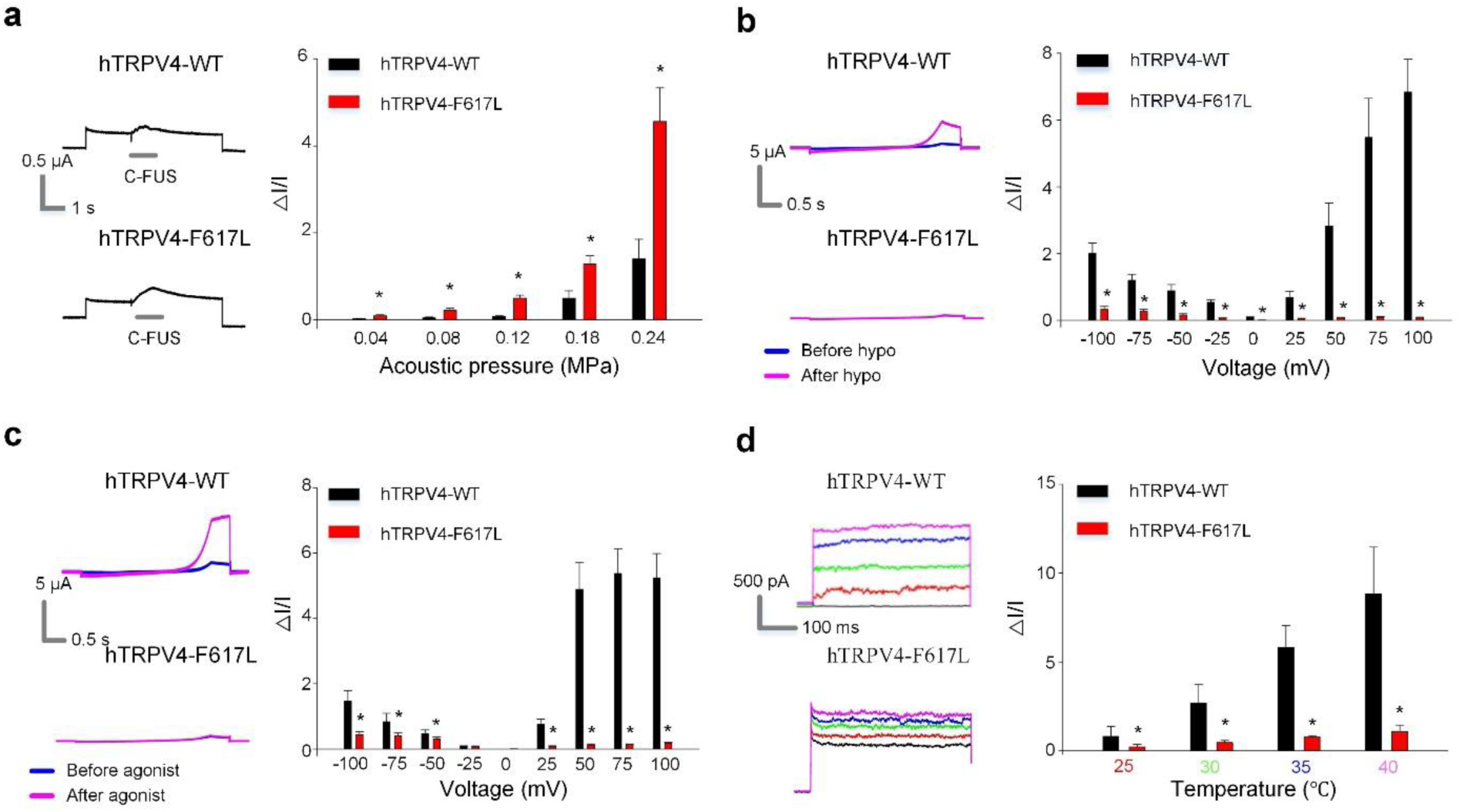
hTRPV4-F617L is more specifically sensitive to FUS. **a-d**, Averaged fold change of current amplitudes of the hTRPV4-WT (black) and hTRPV4-F617L (red) elicited by stimulations of various C-FUS acoustic pressures at a voltage pulse to +50 mV (**Fig. 4b**) (**a**), hypo-osmolarity at various voltages during a voltage ramp (**Fig. S2b**) (**b**), 3 μM 4αPDD at various voltages during a voltage ramp (**Fig. S2b**) (**c**), and various temperatures at a voltage of +80 mV (**Fig. S4c**) (**d**). Sample current traces are shown at the left of each panel.

Ultrasound sensitivity of ion channels has been associated with their mechano- and thermos- sensitivity because ultrasound is capable of altering both the mechanical stress and temperature of the ion channel environment ^18,19,40^. In our experimental conditions (**Fig. S1a**), the heat generated by FUS can be ignored ^18^, and hTRPV4 activation is predominantly caused by the mechanical effects of FUS, supported by experiments with mutations that altered FUS activation differently than activation by elevated temperatures (**Fig. 2**). The mechanosensitive activation of hTRPV4 has been reported previously ^41,42^ but remained a subject of ongoing debate ^43^. This controversy may relate to the observation that whether the channel can be activated by mechanical forces seems to be dependent on how the mechanical forces are applied to the cells ^42^. Our results support the idea that hTRPV4 is indeed a mechanosensitive ion channel, and FUS represents a reliable mechanical stimulation to activate this channel.

Interestingly, the F617L mutation in hTRPV4 is associated with mild metatropic skeletal dysplasia ^44^. Previous studies demonstrated that skeletal dysplasia-associated mutants of hTRPV4 expressed in *Xenopus* oocytes or mammalian cell lines generated larger currents compared to WT channels, suggesting that the disease may be caused by increased basal activity of mutant hTRPV4 leading to enhanced calcium concentrations in chondrocytes ^44,45^. However, among the dysplasia-associated mutations, F617L seems to be exceptional, as it does not increase the basal activity ^45,46^. We compared expression of hTRPV4-WT and hTRPV4-F617L in *Xenopus* oocytes and found no difference in the basal current amplitude within the physiological range of voltages (**Fig. S5a**, **b**). Therefore, unlike other dysplasia-associated mutations, the increased FUS sensitivity suggests that F617L contributes to the disease by increasing hTRPV4 mechano-sensitivity rather than enhancing its basal activity. Accordingly, the enhanced activity of hTRPV4 under mechanical load may cause the increase in chondrocyte intracellular calcium concentration ^46^.

### hTRPV4-F617L structure differs from the hTRPV4-WT channel

To provide an insight into the molecular basis of the hTRPV4-F617L mutation activation, we solved its structure by Cryo-EM (**Fig. 4**, **Fig. S6-8**, **Table S2**). Typical for the vanilloid subfamily of TRP channels, the structure reveals a tetrameric assembly of hTRPV4-F617L subunits, each containing the N-terminal ankyrin repeat domain (ARD), linker domain connecting ARD to the transmembrane domain (TMD), and C-terminal domain which includes an amphipathic TRP helix and a terminal hook. The TMD contains six transmembrane helices, with the first four forming an S1-S4 domain. The last two, S5 and S6 from all subunits, with a re-entrant loop (pore loop or P-loop) between them, assemble an ion conducting channel (**Fig. 4a**). The leucine residue that replaces F617 residues in the middle of S5, at the interface with S6 from the same subunit (**Fig. 4b**). In the hTRPV4-WT, the bulky aromatic side chain of F617 is in the middle of a hydrophobic residue cluster that includes Y602, L614, L618, V620, V708, L710, L711, L714, and Y621, where tyrosine Y621 forms a π-stacking interaction with phenylalanine F617 (**Fig. 4c**). This hydrophobic cluster is right next to the π-bulge in S6, providing an energetic compensation to the energetically unfavorable α-to-π transition in the middle of S6. The hTRPV4-F617L eliminates the bulky aromatic side chain from the center of the hydrophobic cluster and weakens interactions between the contributing residues. This removes the energetic compensation of the α-to-π transition, allowing the network of periodic hydrogen bonds that makes S6 entirely α-helical and straightens it up compared to the kinked S6 in the hTRPV4-WT channel (**Fig. 4b, c**).

**Fig. 4.**
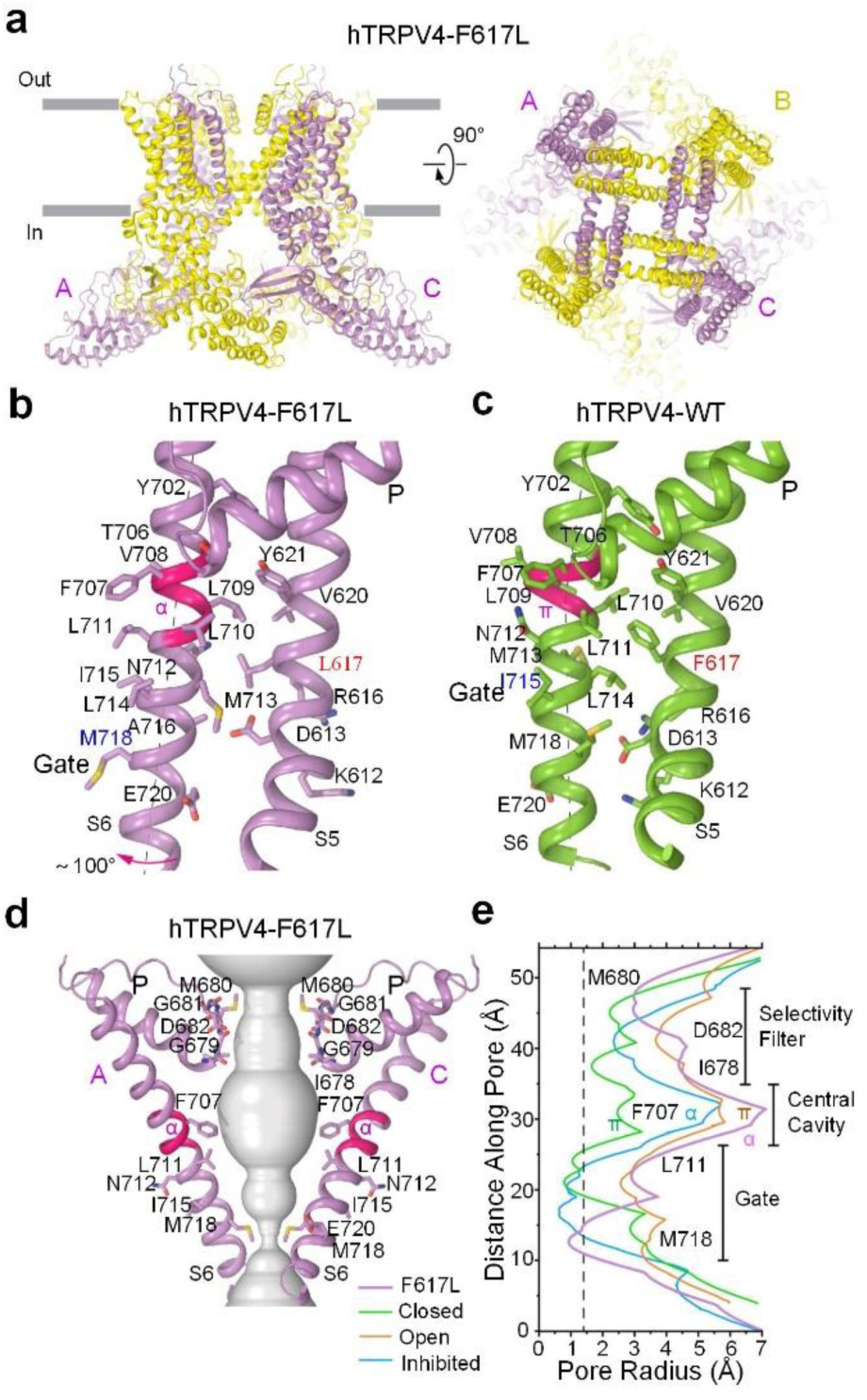
Cyro-EM structure of hTRPV4-F617L. **a**, Side (*left*) and top (*right*) views of hTRPV4- F617L structure. **b**, **c**, Pore-forming S5-S6 region in hTRPV4-F617L (**b**) and hTRPV4-WT (**c**) structures. The segment of S6 undergoing α-to-π transition is colored pink. The side chains of residues contributing to permeation and gating are shown as sticks. **d**, Pore-forming domain in hTRPV4-F617L with the residues contributing to pore lining shown as sticks. Only two of four subunits are shown, with the front and back subunits omitted for clarity. The pore profile is shown as space-filling model (grey). **e**, Pore radius for hTRPV4-F617L, hTRPV4_closed_ (Green; PDB ID: 8T1B), hTRPV4_open_ (Orange; PDB ID: 8T1D), and hTRPV4_inhibited_ (blue; PDB ID: 8T1F) calculated using HOLE. The vertical dashed line denotes the radius of a water molecule, 1.4 Å.

The π-to-α conversion in the middle of S6 in hTRPV4-F617L compared to hTRPV4-WT also results in a ∼100 ° rotation of the C-terminal part of S6, which brings a completely different set of residues to face the ion channel pore. While the pore of hTRPV4-F617L in the apo condition remains apparently closed, it is hydrophobically sealed by the side chains of M718 instead of L711 in hTRPV4-WT. The pore lining in hTRPV4-F617L is also different from the open state of hTRPV4-WT bound to the agonist 4αPDD, which has a wider pore than hTRPV4-WT in the apo state but also contains a π-bulge in the middle of S6. On the other hand, a similar to hTRPV4- F617L pore lining is observed in the antagonist HC067047-bound state of hTRPV4-WT, where S6 undergoes the π-to-α transition ^47^. However, the π-to-α transition in hTRPV4-F617L is also accompanied by a helical-turn elongation of S6 and shortening of the TRP helix (**Fig. S8a-c**). Within the context of one subunit, this transformation results in rotations of the pore domain and ARD relative to the S1-S4 bundle in hTRPV4-F617L compared to any state of hTRPV4-WT. in the context of the tetramer, these rotations cause widening of the TMD and rotation of the intracellular skirt in the hTRPV4-WT open state (**Fig. S8d-f**) ^47^. Similarity of hTRPV4-F617L structure to the open and inhibited states of hTRPV4-WT endows it with features of pre-active and inhibited states. Nevertheless, the hTRPV4-F617L conformation is distinct from any state of hTRPV4-WT revealed before, uncovering the molecular features that allow hTRPV4-F617L to be easily converted to the open state in response to ultrasound but no other stimuli.

### hTRPV4-F617L activation by FUS modulates neural function *in vivo*

We then evaluated the capability of hTRPV4-F617L to sensitize the responsiveness of neurons to ultrasound stimulation in mice. We genetically engineered CaMKⅡ-expressing neurons in the left striatum to express hTRPV4-F617L by directly injecting lentiviruses encoding hTRPV4-F617L under the CaMKⅡ promoter to the left striatum. Control mice were injected with saline. It has been established that activation these neurons can induce rotational motor behavior ^40^. We then tested the activation of those neurons with P-FUS via a freely moving rotational behavior assay (**Fig. 5a**). P-FUS was applied with a central frequency of 3.2-MHz, a pulse repetition frequency (PRF) of 10 Hz, a duty cycle (DC) of 50%, acoustic pressure of 0 and 1.8 MPa, and a burst duration (BD) of 10 s with inter stimulation interval (ISI) of 90 s for a total of five stimulations, as depicted (**Fig. 5b**).

**Fig. 5.**
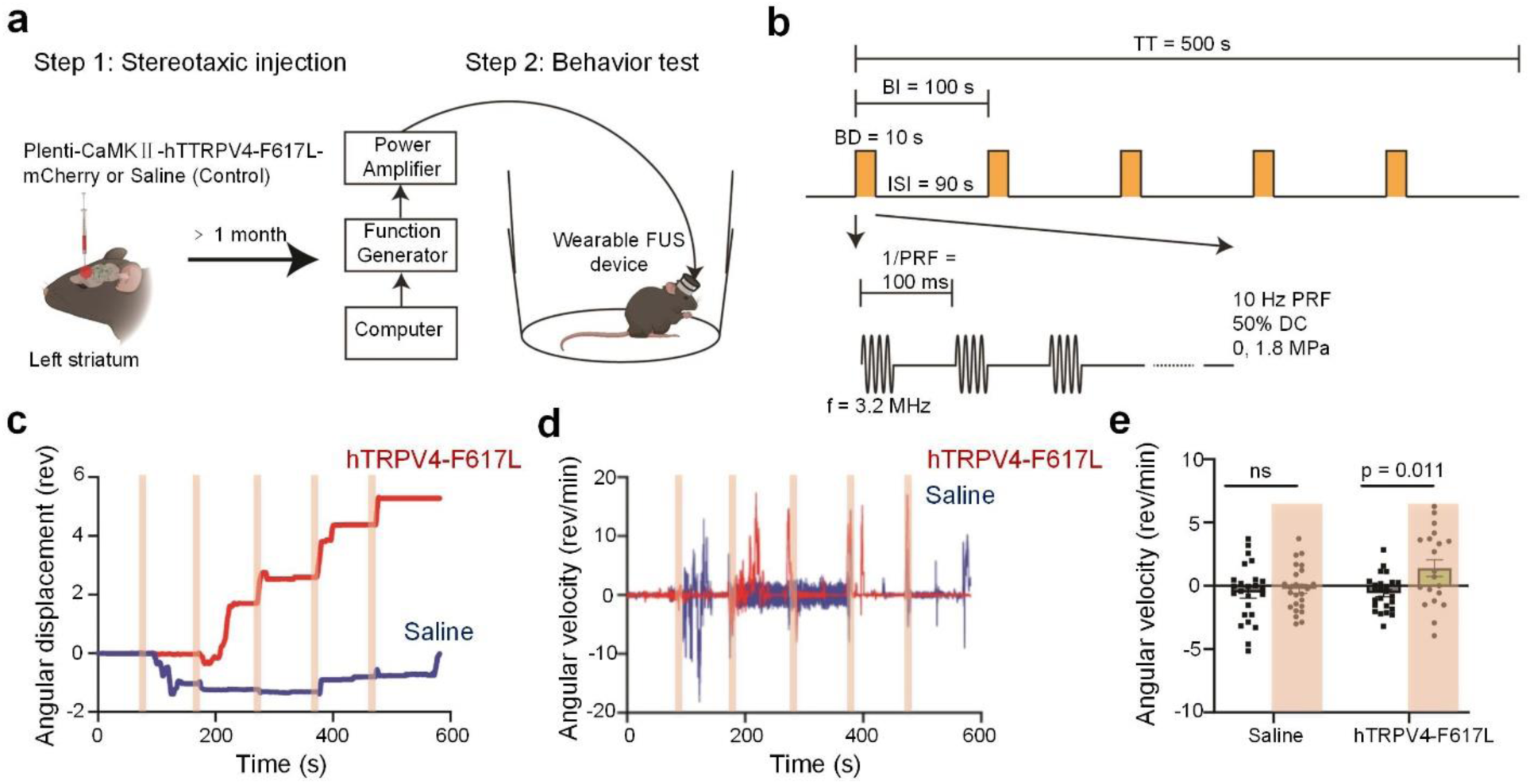
hTRPV4-F617L evokes changes in direction-specific locomotor behavior *in vivo*. **a**, Experimental protocols. **b**, FUS stimulation protocol (TT, total time; BI, burst interval; BD, burst duration). **c**, **d**, Representative plots of the angular displacement and angular velocity over time for a hTRPV4-F617L (red) and a control (blue) mouse. The yellow bars correspond to the application of FUS at an acoustic pressure of 1.8 MPa. **e**, Summary plot of the average angular velocity for control (n = 5) and hTRPV4-F617L (n = 5) at 0 and 1.8 MPa FUS stimulations. Angular velocity values greater than zero correspond to contralateral rotations (clockwise), while angular velocity values less than zero correspond to ipsilateral rotations (counterclockwise). Each point represents on stimulation. Statistical analysis was conducted using two-way repeated measures ANOVA with Bonferroni post-hoc test.

The rotational bias of the mice was assessed by applying ultrasound to the left striatum and examining the angular displacement and angular velocity of the mice (**Fig. 5c, d**). During the P- FUS stimulation, mice expressing hTRPV4-F617L exhibited changes in the angular displacement and angular speed, whereas the control mice showed no significant rotational bias. Group behavior analysis (**Fig. 5e**) further revealed that the angular velocity of hTRPV4-F617L mice during P-FUS stimulation exhibited a statistically significant increase in comparison to the control mice subjected to P-FUS and hTRPV4-F617L mice without P-FUS. These findings confirm that the observed behavioral changes are causally related to hTRPV4-F617L-mediated sonogenetic neuromodulation.

## Conclusion

Our results show that the activation and deactivation of hTRPV4-WT and the mutant hTRPV4- F617L can be precisely controlled by FUS (**Fig. 1, 3**). The hTRPV4-F617L mutant has significantly higher sensitivity to ultrasound than hTRPV4-WT channels (**Fig. 3a**) but minimal sensitivity to other stimuli, including hypotonic stimuli (**Fig. 3b**), agonist 4αPDD (**Fig. 3c**), and elevated temperatures (**Fig. 3d**). The hTRPV4-F617L mutant channel has a normal basal activity, with small open probability at physiological voltages (**Fig. S5a**, **b**), which in combination with the reduction of its sensitivity to stimuli other than FUS minimizes interference with physiological functions when this channel is expressed in neurons. The successful activation of striatal neurons, which causes the rotational behavior in mice (**Fig. 5**), confirms that hTRPV4-F617L can be used as a sonogenetic actuator for *in vivo* cell-type specific manipulations to study neural functions in the brain. The hTRPV4-F617L structure reveals that the F617L mutation causes a significant change in the TMD conformation, the activation gate and the π-bulge in S6 (**Fig. 4**) that plays a critical role in TRP channel activation gating ^47–50^. These results suggest that structure-guided molecular engineering of ion channels is capable of creating more suitable sonogenetic actuators that can be used to understand brain function and develop effective treatments for neurological disorders.

## Cloning and mutagenesis

hTRPV4-WT plasmid was kindly provided by Dr. Hu in Washington University in Saint Louis. To express hTRPV4-WT in *Xenopus* oocytes, the gene was subcloned into a pCDNA 3.1 vector containing a poly A tail. All the point mutations were introduced into hTRPV4-pCDNA 3.1 construct using PCR based site-directed mutagenesis and confirmed it by sequencing the full hTRPV4 insert. cRNA was made by *in vitro* transcription using the mMessage mMachine T7 polymerase kit (Applied Biosystems). To express both hTRPV4-WT and mutants in mammalian cells, the gene was initially placed under the excitatory neuronal promoter kinase Ⅱ alpha subunit (CaMKⅡ). It was then linked with mCherry by the post-transcriptional cleavage linker p2A and finally packaged into a pCDNA 3.1 vector.

For structural studies, cDNA full-length hTRPV4 (NM_021625) was introduced into pEG-BacMam vector for protein expression in mammalian cells ^51^, with C-terminal region coding for the thrombin cleavage site (residues LVPRG), followed by the green fluorescent protein (GFP) and the streptavidin affinity tag (residues WSHPQFEK), as described before ^47^. Introduced hTRPV4-F617L point mutation was confirmed by whole-plasmid sequencing performed by Plasmid saurus using Oxford Nanopore Technology.

## Cell culture and channel expression

*Xenopus* oocytes were collected by laparotomy. All procedures adhere to the recommendations of the Panel on Euthanasia for the American Veterinary Medical Association. Stage Ⅴ or Ⅵ oocytes were isolated by collagenases (0.5 mg/mL, Sigma Aldrich) digestion. hTRPV4-WT or mutant cRNAs (9.2 ng) were micro-injected into each oocyte. The control oocytes that were injected with the same volume of water. The injected oocytes were kept in ND96 solution (in mM): 96 NaCl, 2 KCl, 1.8 CaCl_2_, 1 MgCl_2_, 5 HEPES, 2.5 CH_3_COCO_2_Na, 1:100 Penicillin-streptomycin (Pen-Strep), pH7.6 at 18 ℃ for 2-4 days for electrophysiology recordings (All the chemicals from Sigma Aldrich, and the Pen-Strep from Gibco Life Technology).

Human embryonic kidney 293T (HEK293T) cells were kindly provided by Dr. Chen in Washington University in Saint Louis and cultured in high-glucose Dulbecco’s modified Eagle medium (DMEM, Gibco Life Technologies) supplemented with 10% heat-inactivated fetal bovine serum (FBS, Gibco Life Technologies) and 100 units/mL Pen-Strep (Gibco Life Technologies). The HEK293T cells were incubated at 37 ℃ in a humidified CO_2_ (5%) incubator. Cells were transfected with either hTRPV4-WT or mutant plasmid DNAs (CaMKⅡ-hTRPV4-p2A-mCherry) using the X-tremeGENE^TM^ 9 DNA transfection kit (Sigma Aldrich) 24 h before the experiments. Cells that were transfected by the construct (CaMKⅡ-p2A-mCherry) without hTRPV4 were used as control. The transfected cells were used for both patch clamp and calcium fluorescence imaging.

## Two-electrode voltage clamp recording

Whole-oocyte ionic current following through the ion channels were obtained with the two-electrode voltage clamp. Micro-electrodes were made of glass pipettes (item #B 150-117, Sutter instrument) using a puller (P-1000, Sutter instrument). The two micro-electrodes and the recorded oocyte were immerged in the ND96 bath solution (96 mM NaCl, 2 mM KCl, 1.8 mM CaCl_2_, 1 mM MgCl_2_, 5 mM HEPES, with a pH of 7.6). The GeneClamp 500B amplifier (Axon Instrument) was used to control the voltage clamp. To avoid aliasing, the device applied a low-pass filter to the measured currents at 2 kHz. The ITC-16 board acquired signals at 10 kHz and maintained them using the Patchmaster software (HEKA).

A custom-built two-electrode voltage clamp, incorporating FUS stimulation, was illustrated in **Fig. S1** for conducting ultrasound stimuli experiments. A *Xenopus* oocyte, either expressing hTRPV4 or not, was positioned in the focus region of a chamber that was filled with a bath solution ND96 and subjected to an ultrasound pressure field. A signal-channel arbitrary wave generator (33220A, Agilent) generated a sinusoidal signal that was either continued or pulsed. This signal was then connected to a power amplifier (75A250A, Amplifier Research). The electronic output was directed towards a piezo-ceramic ultrasound transducer (Wuxi Lanhui Electronic) with a focus of 1 mm, an element size of 10 mm, and a central frequency of 1-MHz. The transducer was placed below the recording chamber and submerged in deionized and degassed water. To prevent the production of standing waves in the recording chamber, an absorber was submerged in the recording bath. In this customized designed *in vitro* voltage clamp and FUS stimulation setup, the temperature increase induced by FUS can be ignored because of the brief duration of stimulation time and the presence of a full chamber of ND96 solution (1.5 mL).

The C-FUS and P-FUS were applied during the recording of transmembrane currents. For P-FUS, DC is the ratio of the duration of FUS on to the total duration in one pulse (on and off) in a train of P-FUS. PRF refers to the frequency at which the train of P-FUS is repetitively turned on and off. The ultrasound acoustic pressures were qualified using a lipstick hydrophone (HGL-200, ONDA Corporation) placed in a water tank filled with deionized and degassed water at a room temperature, and reported acoustic pressures were based on these measurements.

For hypotonic stimuli experiments, a hypotonic solution (composed of 66 mM KCl, 1.8 mM BaCl_2_ and 10 mM K^+^-HEPES with pH of 7.4, without sorbitol) was perfused onto oocytes to steady state. The current comparison of hTRPV4-WT between ND96 solution of iso/hypotonic solution is depicted in **Fig. S9**. In the experiments shown in **Fig S5c, d** the above isotonic and hypotonic solutions were used. However, in the experiments shown in **Fig S5e-h**, a different isotonic solution (in mM): 105 NaCl, 6 CsCl, 5 CaCl_2_, 1 MgCl_2_, 90 D-Manitol, 10 glucose, 10 HEPES, pH7.4 and hypotonic solution (composition the same as isotonic solution but without D-Manitol) were used (All the chemicals are from Sigma Aldrich). For agonist stimuli experiments, 4αPDD (Sigma Aldrich) were stored as 3 mM stocks in dimethyl sulfoxide (DMSO) and diluted in bath solutions to working concentration each experiment day.

## Calcium fluorescence imaging

A customized experimental setup was established to enable a simultaneous monitoring of fluorescence and the application of FUS stimulation to HEK293T cells (**Fig. S2c**). The cells that underwent transfection were loaded with the fluorescence Ca^2+^ indicator Fluo-4 AM (Thermo Fisher Scientific) according to the manufacturer’s instructions. The fluorescent microscope (LX70, Olympus) was used to capture images of the dynamic Ca^2+^ response to FUS stimulation. During calcium imaging, the transfected cells were placed in a live cell imaging solution (Thermo Fisher Scientific) in the presence of 10 mM glucose (Thermo Fisher Scientific). The expression of hTRPV4 was confirmed by the observation of Ca^2+^ influx in response to 4αPDD (3 μM) (**Fig. S2a**, **b**). A FUS with a central frequency of 1.7 MHz, a peak negative pressure of 0.568 MPa, a DC of 40%, and a PRF of 10 Hz was applied to the transfected cells for a duration of 5 s.

## Patch clamp recording

Patch clamp pipettes were fabricated by pulling borosilicate glass tubes (Sutter Instrument) using a P-97 micro-pipette puller (Sutter Instrument) and then fire-polished to achieve resistance ranges from 2 to 6 MΩ for microscopic recordings. A Patchmaster software (HEKA) controlled an EPC 10 USB patch clamp amplifier. The current signal was sampled at a frequency of 10 kHz and filtered at a frequency of 2.25 kHz. The holding membrane potential was set at 0 mV, followed by a depolarizing step to +80 mV and hyperpolarizing step to -80 mV. The temperature of the bath solution was controlled using a SHM-828 eight-line heater operated by a CL-100 temperature controller (Warm Instruments). The transmembrane of the solution was measured using a BAT-12 microprobe thermometer (Physitemp Instrument). The thermos-probe was positioned within a proximity of 1 mm from the cell. The patch clamp recording utilized symmetrical bath and pipette solution. These solutions consisted of 160 mM NaCl, 2 mM CaCl_2_, 1 mM MgCl_2_, and 10 mM HEPES (pH7.4) (All chemicals are from Sigma Aldrich).

## Protein expression and purification

hTRPV4 bacmids and baculoviruses were produced using standard procedures ^51^. Briefly, baculovirus was made in Sf9 cells (Thermo Fisher Scientific, mycoplasma test negative, Gibco #12659017) for ∼120 hours and added to suspension-adapted HEK293T cells lacking N-acetyl-glucosaminyl transferase I (GnTI^-^, mycoplasma test negative, ATCC #CRL-3022) that were maintained at 37 ℃ and 5% CO_2_ in Freestyle 293 media (Gibco Life Technology) supplemented with 2% FBS (Gibco Life Technology). To reduce hTRPV4 cytotoxicity, 10 μM ruthenium red (Thermo Fisher Scientific) was added to the suspension of HEK293T cells. To enhance protein expression, sodium butyrate (10 mM, Thermo Fisher Scientific) was added 24 hours after transfection, and the temperature was reduced to 30 ℃. The cells were harvested 72 hours after transfection by 15 min centrifugation was 5,471 g using a Sorvall Evolution RC centrifuge (Thermo Fisher Scientific). The cells were washed in the phosphate buffer saline (PBS, pH8.0) and pelleted by centrifugation at 3,202 g for 10 min using an Eppendorf 5810 centrifuge.

hTRPV4 was purified based on our previous established protocols ^47^. In a nutshell, the cell pellet was re-suspended in the ice-cold buffer containing 10 mM Tris (pH8.0), 150 mM NaCl, 0.8 μM aprotinin, 4.3 μM leupeptin, 2 μM pepstatin A, 1 μM phenylmethylsulfonyl fluoride (PMSF), and 1 mM β-mercaptoethanol (β-ME) (All chemicals are from Sigma Aldrich). The suspension was supplemented with 1% (w/v) glycol-diosgenin (GDN, Sigma Aldrich), and cells were lysed at constant stirring for 1 hour at 4 ℃. Unbroken cells and cell debris were pelleted in the Eppendorf 5810 centrifuge at 3,202 g and 4 ℃ for 10 min. Insoluble materials was removed by ultra- centrifugation for 1 hour at 186,000 g in a Beckman Coulter centrifuge using a 45 Ti rotor. The supernatant was added to the strep resin, which was then rotated for 20 min at 4 ℃. The resin was washed with 10 column volumes of wash buffer containing 10 mM Tris (pH8.0), 150 mM NaCl, 1 mM β-ME, and 0.01 (w/v) GDN, and the protein was eluted with the same buffer supplemented with 2.5 mM D-desthiobiotin (Sigma Aldrich). The eluted protein was concentrated to 0.5 mL using a 100-kDa NMWL centrifugal filter (Millipore Sigma^TM^, Amicon^TM^) and then centrifuged in a Sorvall MTX 150 Micro-Ultracentrifuge (Thermo Fisher Scientific) for 30 min at 66,000 g and 4 ℃ using a S100AT4 rotor before injecting it into a size-exclusion chromatography (SEC) column. The protein was purified using a Superose TM 6 10/300 GL SEC column attached to an AKTA FPLC (GE Healthcare) and equilibrated with the buffer containing 150 mM NaCl, 20 mM Tris (pH8.0), 1 mM β-ME, and 0.01 (w/v) GDN. The tetrameric peak fractions were pooled and concentrated to 3.2 mg/mL using a 100-kDa NMWL centrifugal filter (Millipore Sigma ^TM^, Amicon^TM^).

## Cyro-EM sample preparation and data collection

Prior to sample application, UltrAuFoil R 1.2/1.3 (Au300) grids were plasma treated in a PELCO easiGlow glow discharge cleaning system (0.26 mBar, 15 mA, “glow” for 20 s, and “hold” for 10 s). A Mark Ⅳ Vitrobot (Thermo Fisher Scientific) set to 100% humidity at 4 ℃ was used to plunge-freeze the grids in liquid ethane after applying 3 μL of protein sample to their gold-coated side using the blot time of 3 s, blot fore of 3 s, and wait time of 20 s. The grids were stored in liquid nitrogen before imaging.

## Imaging processing and 3D reconstruction

Cryo-EM data were processed in cryoSPARC 4.4.0 ^52^. 14,100 movie stacks were aligned using Patch Motion Correction algorithm and then subjected to patch CTF estimation. The obtained micrographs were semi-manually curated based on their relative ice thickness and CTF fit resolution, with micrographs that had a predicted resolution high than 6 Å being excluded. A total number of 2,170,956 particles were picked using blob picking, the best-resolved 2D classes were used as templates for template picking. These particles extracted using a 320-pixel box size and subsequently binned to a 160-pixel box size for cleaning through several rounds of 2D classification and heterogeneous refinement. A final subset of 605,150 particles was re-extracted with the 320-pixel box size and used for non-uniform refinement with C2-symmetry and global and local CTF refinement. To obtain a more homogeneous fraction of particles, they were classified by ab-initio reconstruction with 5 classes, from which the largest class was taken for non-uniform refinement and additional round of ab-initio reconstruction with 4 classes. Ab-initio reconstruction results were taken to heterogeneous refinement. Non-uniform refinement of the largest class (178,505 particles) gave a 3.74 Å reconstruction (*C1*). To improve quality of the reconstruction, two local refinements were done on this subset of particles: with a focused mask covering hTRPV4-F617L and excluding RhoA and with a focused mask around TMD (**Fig S7**). 3.66 Å reconstruction (*C2*) of full-length hTRPV4-F617L and 4.10 Å reconstruction (*C2*) of the TMD were united into a composite map by the combine focused maps algorithm implemented in Phenix.

## Model building

The hTRPV4-F617L model was built in Coot ^53^ using the previous published cryo-EM structure of hTRPV4 (PDB ID: 8T1B) as a guide. The model was tested for overfitting by shifting their coordinates by 0.5 Å (using Shake) in Phenix ^54^, refining the shaken model against the corresponding unfiltered half map, and generating densities from the resulting model in UCSF ChimeraX. The resulting model was real space refined in Phenix 1.18 and visualized using UCSF ChimeraX, and PyMOL (The PyMOL Molecular Graphics System). The pore radius was calculated using HOLE ^55^.

## *In vivo* neuromodulation

Sonogenetic neuromodulation was performed following our previous procedures ^40^. Mice (C57BL/6 mice, 6-8 weeks old) were intracranially injected with lentiviruses encoding hTRPV4- F617L under the CaMKⅡ-expressing neurons to expressing hTRPV4-F617L. A total of 1250 μL of hTRPV4-F617L virus (4.8 × 10^8^ viral particles/mL) was injected at a rate of 64 nL/min to the left striatum. Control mice were injected with 1250 nL saline to match the injection volume delivered to the striatum. Cloning, packaging, purification, and viral titer calculations were performed by the Hope Center Viral Vectors Core at Washington University School of Medicine.

A wearable FUS device was used to stimulate the striatum of freely moving mice. The FUS device operated at a frequency of 3.2-MHz with a 13 mm aperture and a 10 mm radius of curvature. The wearable FUS transducer was plugged into a 3D printed circular adapter attached to the mouse skull, ensuring alignment of the FUS focus with the left striatum. Ultrasound gel was applied to couple the transducer to the adaptor for optimal transmission.

Following a 2-day adaption period to the behavior testing environment, the locomotor behavior of the mice in response to FUS stimulation was assessed. FUS was applied at a DC of 50%, PRF of 10 Hz, and 10 s total sonication duration with 90 s inter-stimulation interval for a total of 5 stimulations. The onset and offset of the ultrasound pulse were smoothed to avoid possible auditory effects ^56^. The acoustic pressures used in the study were 0 and 1.8 MPa (in situ with skull attenuation considered). Custom MATLAB software was used to control ultrasound application via an Arduino Uno. A light on the Arduino Uno would turn on when ultrasound was applied to precisely synchronize mouse behavior to each FUS stimulation. Mouse behavior was recorded using a video camera (Logitech C920X, 30 fps) before, during and after each focused ultrasound stimulation. All recorded videos were processed using Bonsai to qualify the positional coordinates and the angular orientation of the mice. These data were then analyzed using a custom MATLAB script to compute the average angular velocities.

## Statistics

All the figures presented in the results as average ± standard error of the mean (SEM). Statistical differences were considered significant whenever p < 0.005.

---

**Supplementary Figure 1.**
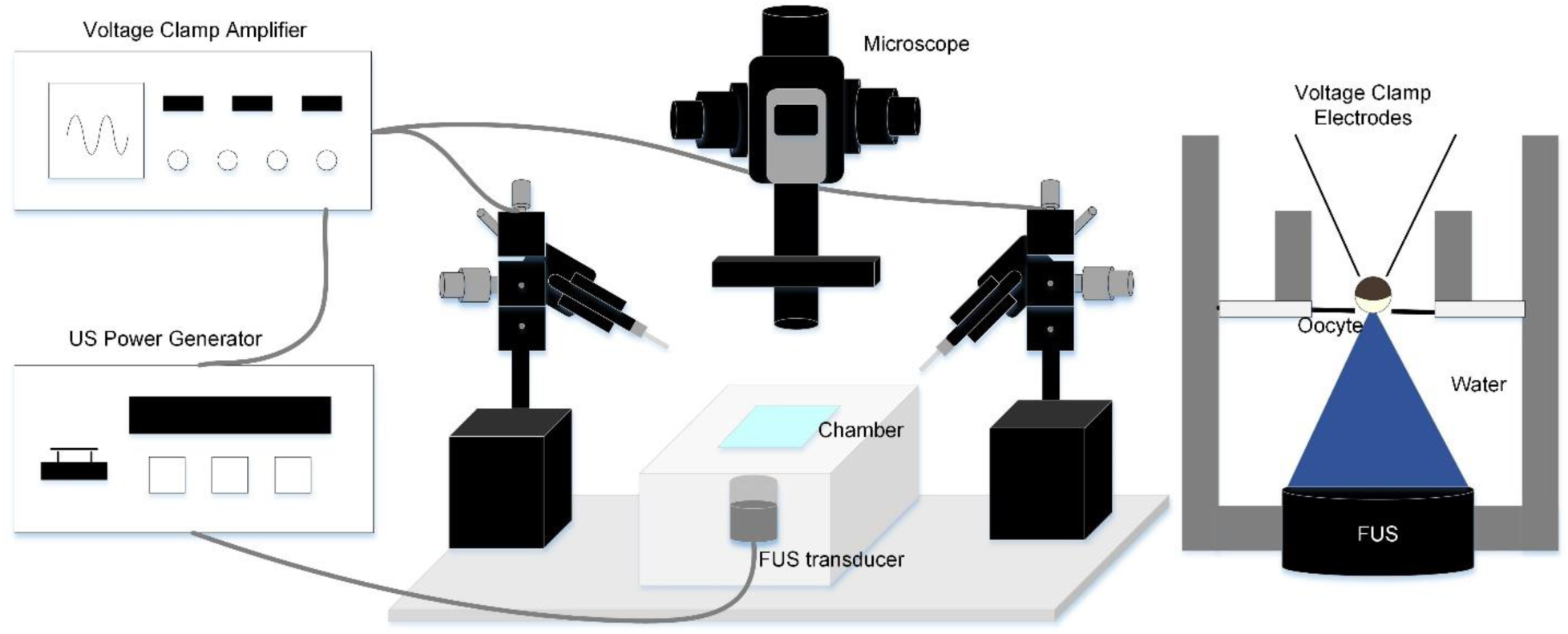
The FUS-two-electrode voltage clamp recording system. Left shows the instrument set up and connections. The recording chamber (indicated in cyan) is enlarged and shown in the right. This transducer with a focus of approximately 10 mm^3^ covered the entire size of the oocyte with a typically diameter about 0.8 mm.

**Supplementary Figure 2.**
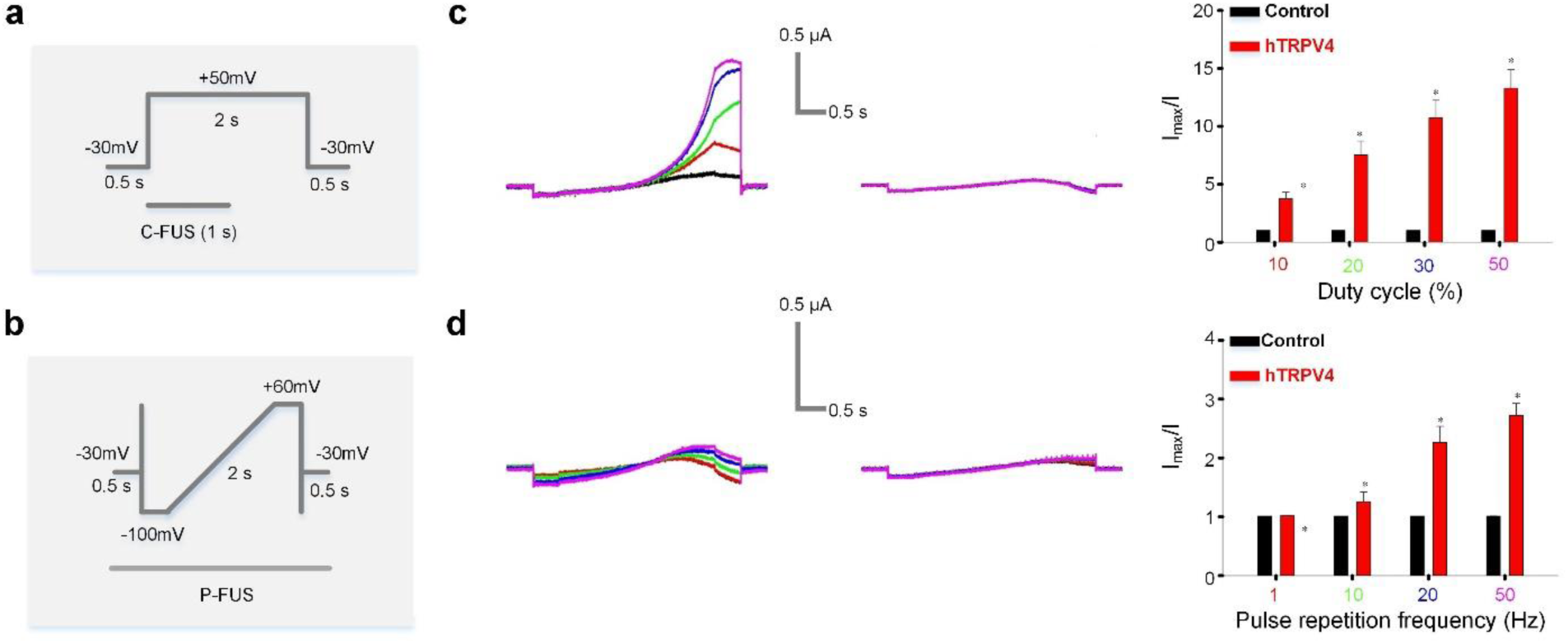
Voltage protocols for eliciting hTRPV4 currents and P-FUS activation. **a**, Voltage protocols used in **Fig. 1a**. **b**, Voltage protocols used in **Fig. 1b-c**, **2b-c**, **S4e- h**, **S5c, e**, **S9a**, and **c-d** in this figure. **c, d**, Activation of hTRPV4-WT by P-FUS with 0.12 MPa, 10 Hz PRF, and various DC (**c**) and 0.06 MPa, 10% DC, and various PRF (**d**). Currents of a hTRPV4-WT-expressed oocyte (*left*) and a water-injected control oocyte (*middle*) are shown. Colors indicate the P-FUS parameters as in (*right*), the averaged fold change of current amplitudes elicited by P-FUS at various parameters. The results in **c** suggest that an increased P-FUS exposure in duration enhanced currents in oocytes expressing hTRPV4-WT. Interestingly, an increase of PRF exposure with a constant P-FUS acoustic pressure and DC also resulted in a rise of hTRPV4 currents. **d**, The PRF increase did not alter the total FUS exposure of the oocytes expressing hTRPV4-WT, while reduced the internal among trains of P-FUS stimulations. This result suggests an accumulative effects of FUS stimulation of hTRPV4 activation in time.

**Supplementary Figure 3.**
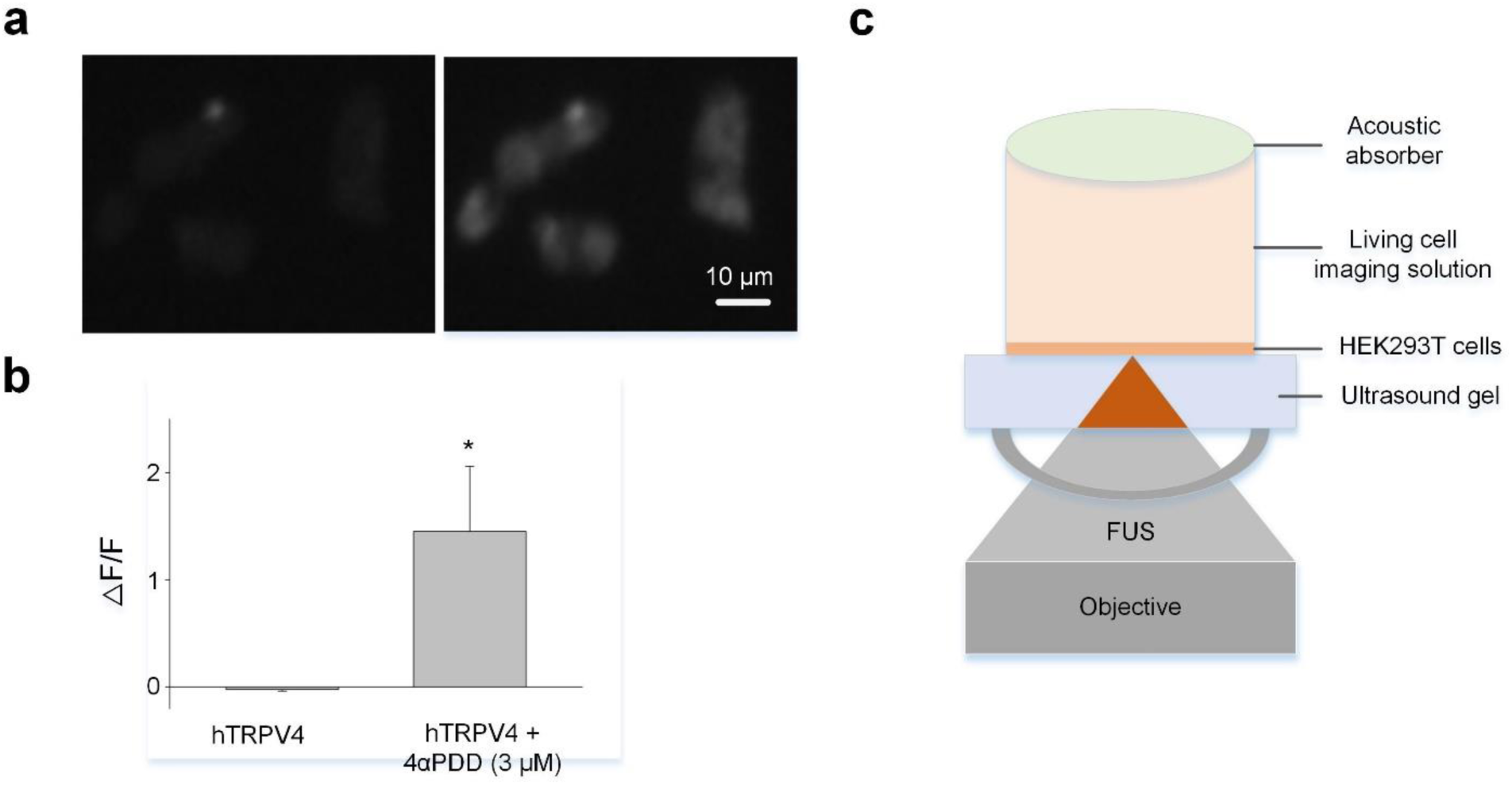
Recording fluorescence of HEK293T cells. **a**, The effects of 4αPDD on hTRPV4-expressed HEK293T cells. The HEK293T cells were transfected with hTRPV4 (plenti-CaMKⅡ-hTRPV4-p2A-mCherry) plasmids and located with Ca^2+^ fluorescence indicator, Fluo-4 AM. Image before (*left*) and after (*right*) adding 4αPDD (3 μM). **b**, Normalized fluorescence change (△F/F) in hTRPV4-expressing cells with or without 4αPDD. **c**, Illustration of the experimental setup for *in vitro* simultaneous FUS stimulation and Ca^2+^ fluorescence imaging. The placement of the ring-shaped transducer was adjusted to achieve confocal alignment with the objective lens of the fluorescence microscope. For an optimal alignment directly above the ring-shaped transducer, a transparent plastic window measuring 0.2 mm in thickness was designed, accompanied by a thin layer of degassed ultrasound coupling gel. The well was filled with live cell imaging solution. An ultrasound absorber was placed on top of the well to reduce the presence of the standing wave.

**Supplementary Figure 4.**
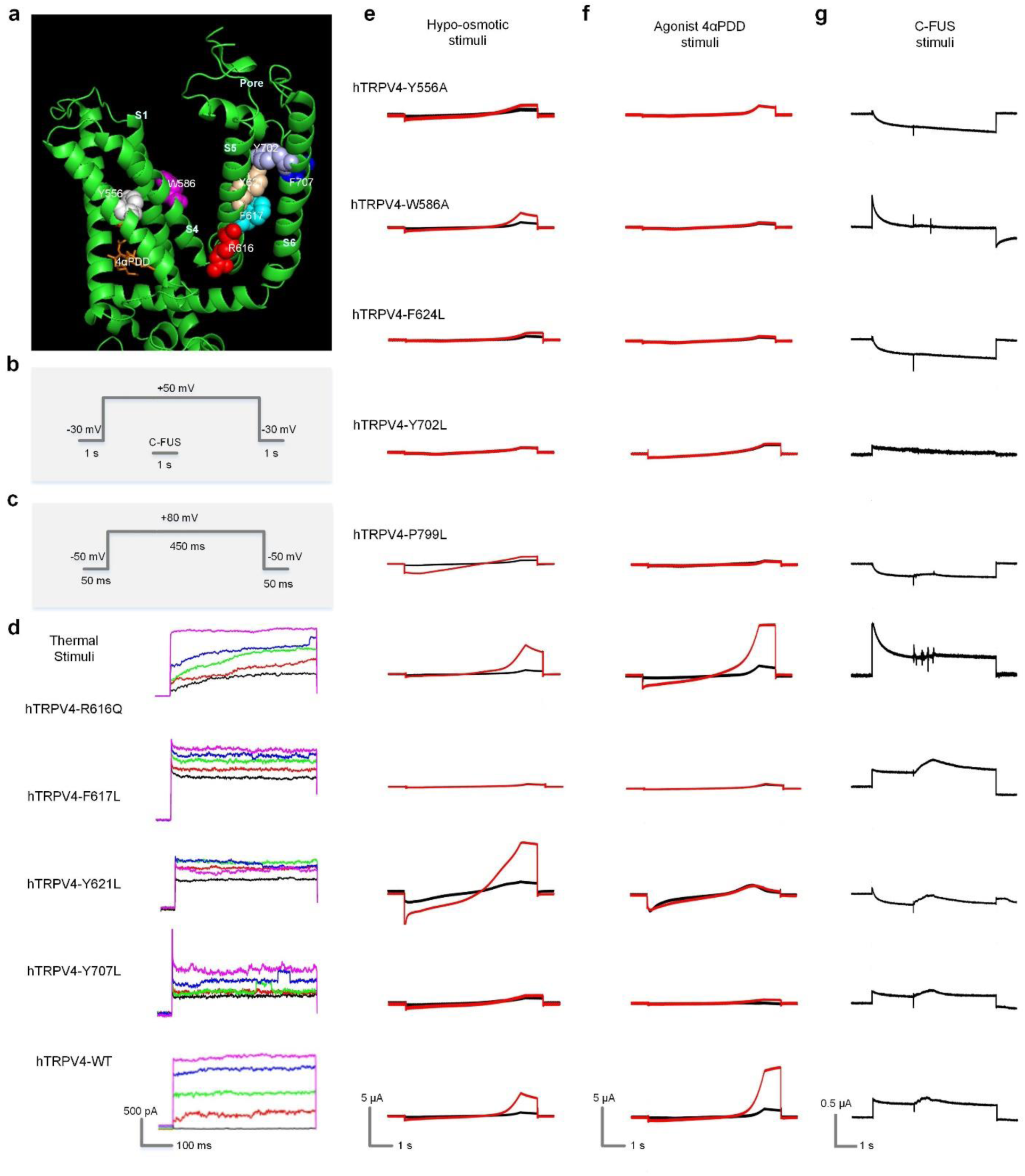
Measuring sensitivity of mutant hTRPV4 to ultrasound and other stimuli. **a**, Mapping the mutated amino acids on the Cryo-EM structure of hTRPV4 in complex with 4αPDD (PDB ID: 8T1D). **b, c**, Voltage protocols employed for C-FUS stimulation in **g** (**b**) F707L, Y556A, W586A, F624L, Y702L, and P799L) and hTRPV4-WT (labeled at the left most current traces) activated by temperature (**d**), hypo-osmolarity (**e**), 4αPDD (**f**), and C-FUS (**g**). In d, the colors indicate different temperatures (℃): black, 20; red, 25; green, 30; blue, 35; magenta, 40. In **e** and **f**, the currents were elicited by a voltage ramp (**Fig. S2b**). The black and red traces are currents before and after stimulation of hypo-osmolarity (**e**) and 3 μM 4αPDD (**f**). In **g**, C-FUS had 1-MHz central frequency and 0.18 MPa for a duration of 1 s.

**Supplementary Figure 5.**
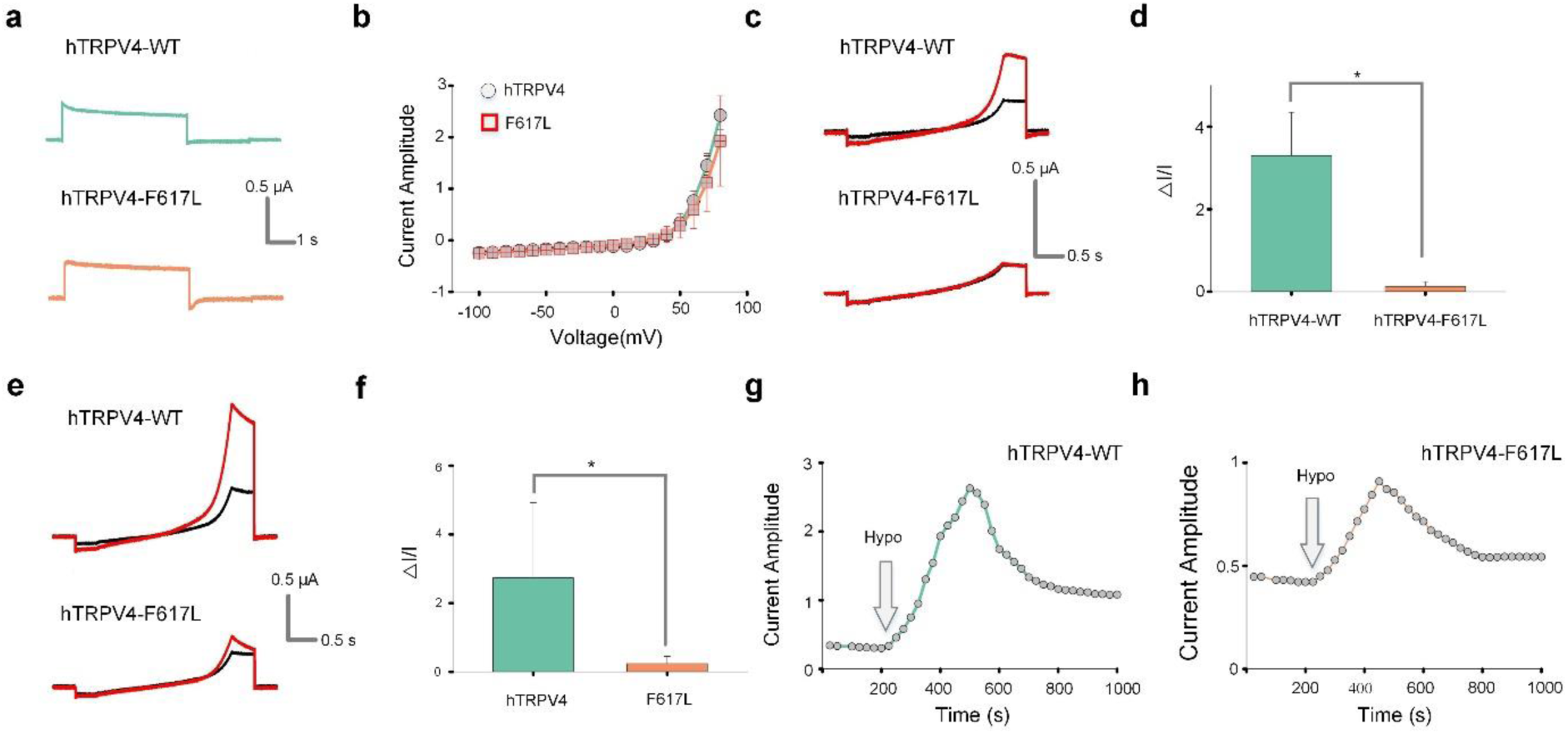
The effects of mutation F617L on hTRPV4 expression and sensitivity to hypo-osmolarity stimulation. Previous studies showed the results hTRPV4-F617L either reduced ^38^ or did not significantly alter the expression level ^45^ of the expressed hTRPV4 currents. Our experimental results show that the hTRPV4-WT and the hTRPV4-F617L have similar current expression at all tested voltages (**a, b**). On the other hand, previous reports showed conflicting results on whether hTRPV4-F617L decreased ^45^ or enhanced ^38^ the hypo-osmolarity stimulation of hTRPV4 activation. we used the isotonic/hypotonic solutions described in the former ^30,45^ (**c, d**) and latter ^38^ (**e-h**) reports, respectively, to test the effects of hTRPV4-F617L on hypo-osmolarity stimulation of hTRPV4 activation. In both conditions, hTRPV4-F617L reduced hypo-osmolarity stimulation consistently. **a, b**, The hTRPV4-WT and hTRPV4-F617L currents expressed in oocytes studied in parallel. Sample current traces elicited by a voltage pulse at +40 mV from a holding potential of -30 mV (**a**) and averaged peak current amplitudes at different voltages (**b**). **c**, Stimulation of hTRPV4-WT and hTRPV4-F617L currents by hypotonic solution (red) from isotonic solution (black). The currents were elicited by a voltage ramp from -100 mV to +60 mV. The isotonic/hypotonic solution had the composition as previously described ^30,45^. **d**, The normalized peak current change caused by hypotonic solution as shown in (**c**). **e, f**, Similar with (**c, d**) except that the isotonic/hypotonic solutions had the composition as described by a different report ^38^. **g, h**, Time course of current change after the replacing of isotonic solution with shown for hypo-osmolarity stimulation in this study were recorded after the current reached the steady state. n > 6 for all experiments shown in this figure.

**Supplementary Figure 6.**
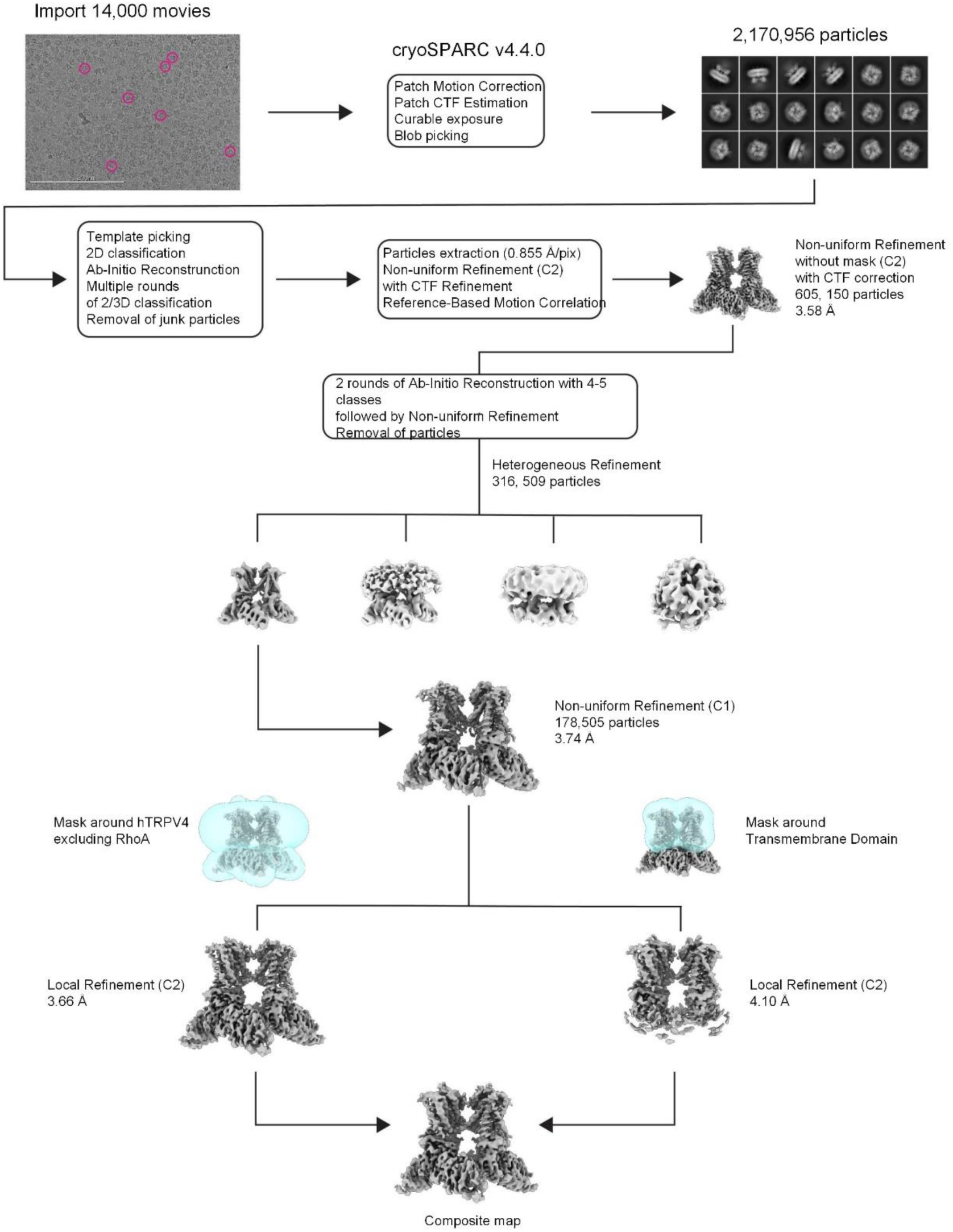
Cyro-EM data processing workflow for hTRPV4-F617L. On the top, a representative of the 14,000 collected micrographs with example particles circled in pink and 2D class average.

**Supplementary Figure 7.**
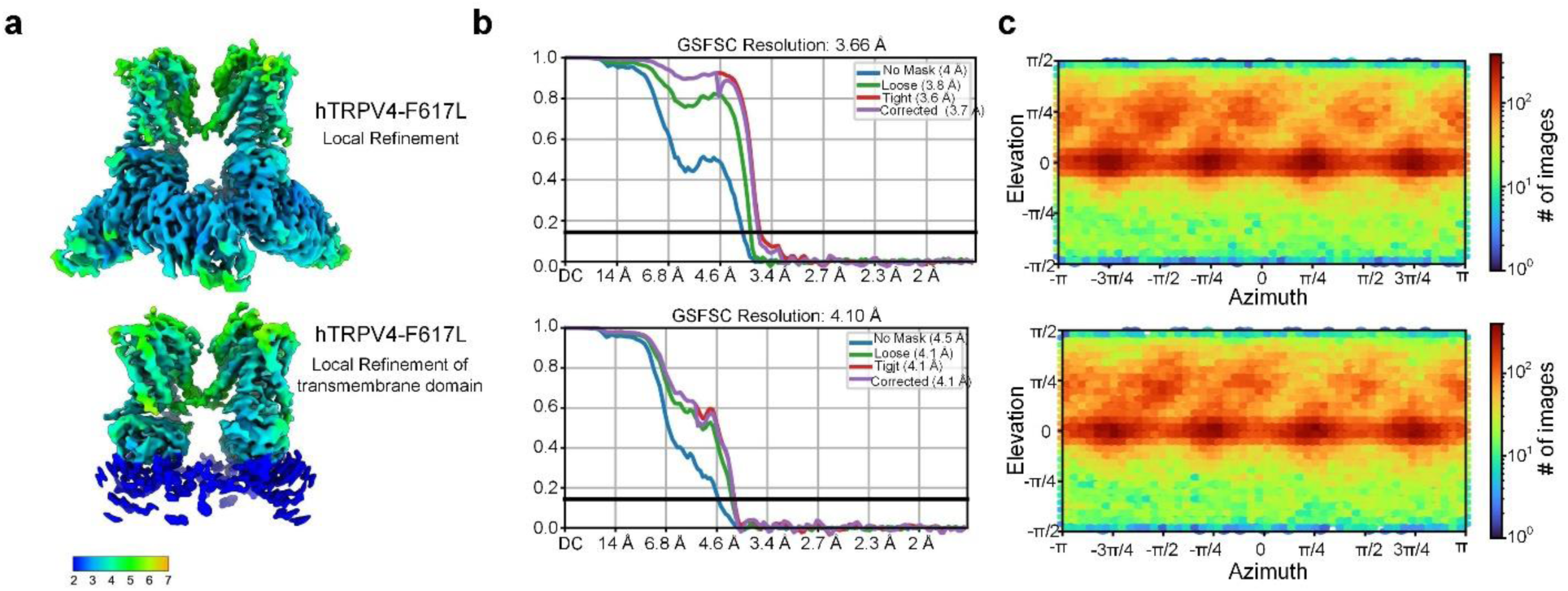
Characteristics of hTRPV4-F617L Cryo-EM reconstructions. Cryo-EM maps colored according to the local resolution estimated by cryoSPARC with color scale in Å (**a**), FSC curves calculated between half maps, with the overall resolution estimated using the FSC = 0.143 criterion (**b**), and angular distribution of particles calculated using the 3D refinement reconstruction algorithm in cryoSPARC (**c**) are shown for the local refinement of full length hTRPV4-F617L (*upper row*) and the local refinement of the TMD (*lower row*).

**Supplementary Figure 8.**
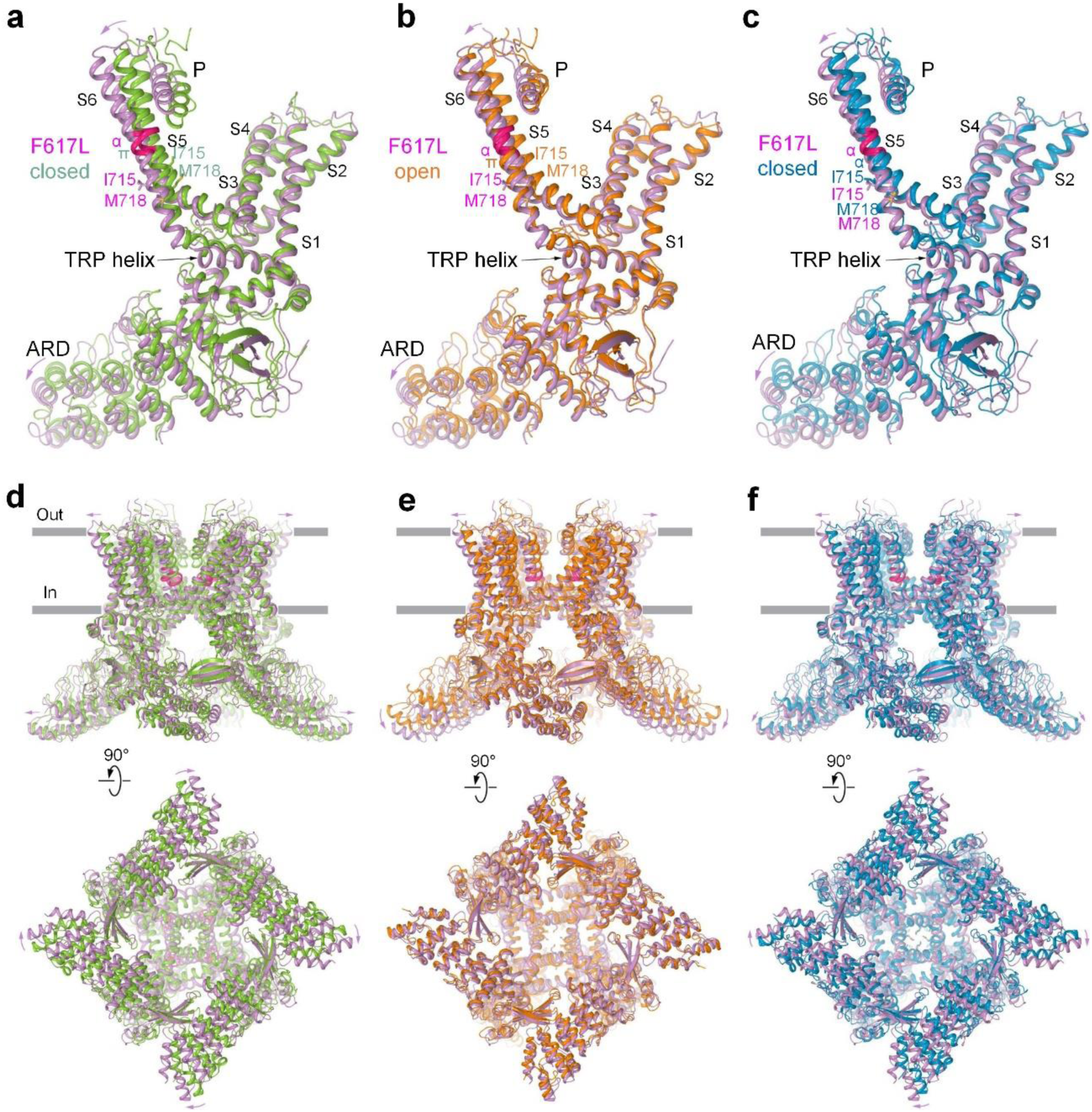
Comparison of hTRPV4-F617L and hTRPV4-WT structures. **a-c**, S1-S4-based superposition of a single hTRPV4-F617L subunit (violet) with hTRPV4-WT subunit in the closed (**a**, green; PDB ID: 8T1B), open (**b**, orange; PDB ID: 8T1D), and inhibited (**c**, blue; PDB ID: 8T1F) states. The region undergoing α-to-π transition is colored pink. The side chains of residues contributing to the gate are shown as sticks. **d-f**, S1-S4-based superposition of the hTRPV4-F617L tetramer (violet) with the tetramer of the closed (**d**, green; PDB ID: 8T1B), open (**e**, orange; PDB ID: 8T1D), and inhibited (**f**, blue; PDB ID: 8T1F) state structures of hTRPV4- WT. Violet arrows illustrate relative movement of domains.

**Supplementary Figure 9.**
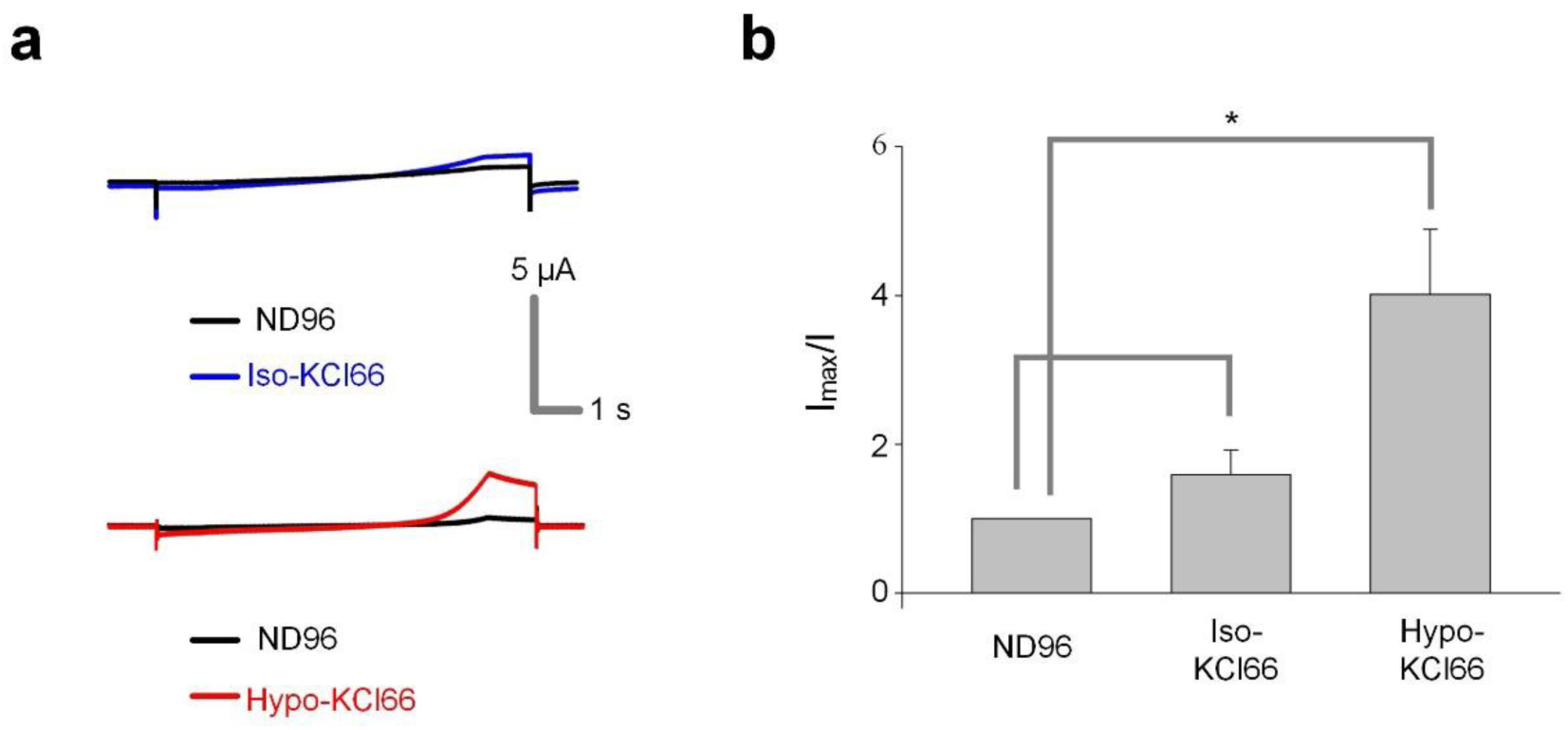
hTRPV4-WT currents in ND96 and Iso/Hypo KCl66 solution. **a**, Currents of hTRPV4-WT in ND96 (black) as compared in Iso-KCl66 (blue, *upper*) and Hypo- KCl66 (red, *lower*) solutions. The currents were elicited by a ramp voltage protocol (**Fig. S2b**). **b**, Averaged fold change of current amplitude of the hTRPV4-WT in ND96, Iso-KCl66, and Hypo- KCl66 solutions, respectively. *: p < 0.005 (n > 5).

**Supplementary Table 1.** Summary of the sensitivity of hTRPV4-WT and hTRPV4 mutants to various stimuli. “✔” means yes; “**↓**” means decreased sensitivity; “**↑**” means increased sensitivity; “NS” means no significant difference.

| hTRPV4 | Ultrasound sensitivity | Hypo-osmotic sensitivity | Agonist 4αPDD sensitivity | Temperature sensitivity |
| --- | --- | --- | --- | --- |
| WT | ✓ | ✓ | ✓ | ✓ |
| Y556A | ↓ | ↓ | ↓ |  |
| W586A | ↓ | NS | ↓ |  |
| R616Q | NS | ↑ | NS | NS |
| F617L | ↑ | ↓ | ↓ | ↓ |
| Y621L | NS | ↑ | ↓ | ↓ |
| F624L | ↓ | ↓ | ↓ |  |
| Y702L | ↓ | ↓ | ↓ |  |
| F707L | NS | ↓ | ↓ | ↓ |
| P799L | NS | NS | ↑ |  |

**Supplementary Table 2.** Cryo-EM data collection, refinement and validation statistics.

|  | Full-length TRPV4 F617L<br>Apo<br>(EMDB-xxxx)<br>(PDB xxxx) |
| --- | --- |
| **Data collection and processing** |  |
| Magnification | 105,000x |
| Voltage (kV) | 300 |
| Electron exposure (e-/Å<sup>2</sup>) | 50.8 |
| Defocus range (µm) | -0.75 to -1.75 |
| Reported pixel size (Å) | 0.855 |
| Calibrated pixel size (Å) | 0.855 |
| Exposures (no.) | 14,100 |
| **Processing software** |  |
| Platform software for particle picking | CryoSPARC v4.4.0 |
| Motion correction | MotionCor2 |
| CTF estimation | Patch CTF |
| Software for 2D/3D class. & refinements | Cryosparc v4.4.0 |
| Symmetry imposed | C4 |
| Initial particle images (no.) | 2,170,956 |
| Final particle images (no.) | 178,505 |
| Map resolution (Å) | 3.66 |
| FSC 0.143 |  |
| Map resolution range (Å)<br>min/median/max | 2.15/5.26/42.55 |
| **Refinement** |  |
| Initial models used (PDB code) | 8T1B |
| Model resolution (Å) | 4.10 |
| FSC threshold |  |
| Map sharpening *B* factor (Å<sup>2</sup>) | -99.5 |
| Model composition |  |
| Non-hydrogen atoms | 20,584 |
| Protein residues | 2,564 |
| Ligand atoms | 0 |
| R.m.s. deviations |  |
| Bond lengths (Å) | 0.007 |
| Bond angles (°) | 1.263 |
| Validation |  |
| CC (across whole map volume) | 0.6491 |
| CC (only across atoms in the model) | 0.7440 |
| Clashscore, all atoms | 4.25 |
| Poor rotamers (%) | 0.53 |
| Ramachandran plot |  |
| Favored (%) | 84.50 |
| Allowed (%) | 14.72 |
| Disallowed (%) | 0.78 |

## Acknowledgements

This work was supported by the National Institute of Health (NIH) grant R01MH116981 and R01 NS12846 for HC; R21HL161629 for JC and HC; R37NS083660, R01NS107253, R01AR078814, and R01CA206573 for AIS; R01NS03954 for JZ. This work was also supported by the McDonnell Center for Cellular and Molecular Neurobiology of Washington University in Saint Louis-FY21 Post-Doctoral Fellowship for LZ and the Hope Center Viral Vectors Core at Washington University in Saint Louis.

## Author Contribution

LZ, JS, and JC performed mutagenesis and voltage clamp studies, LZ, KX, YY and SZ performed experiments to record FUS activation of hTRPV4 in HEK293T cells. KX, YY, SZ, and HC performed *in vivo* neuromodulation studies. IT and AIS performed structural studies using the cryo-EM approach. SL and JZ performed patch clamp studies of WT and mutant hTRPV4 channels dependence on temperature. LZ, KX, IT, AIS, HC, and JC wrote the manuscript with the input from all authors.

## Additional Information

Supplementary Information is available for this paper. Correspondence and requests for materials should be addressed to Lu Zhao, Jianmin Cui, Hong Chen, and Alexander I. Sobolevsky. Reprints and permissions information is available at www.nature.com/reprints.

